# Triple oral beta-lactam containing therapy for Buruli ulcer treatment shortening

**DOI:** 10.1101/439828

**Authors:** María Pilar Arenaz Callao, Rubén González del Río, Ainhoa Lucía Quintana, Charles J. Thompson, Alfonso Mendoza-Losana, Santiago Ramón-García

**Author notes:** Corresponding authors. Mailing addresses: Department of Microbiology, Preventive Medicine and Public Health. Faculty of Medicine. University of Zaragoza. C/ Domingo Miral s/n. 50009. Zaragoza, Spain. Global Health R&D, GlaxoSmithKline, C/ Severo Ochoa 2, 28760, Tres Cantos, Madrid, Spain. mail.

## Abstract

The potential use of clinically approved beta-lactams for Buruli ulcer (BU) treatment was investigated with representative classes analyzed *in vitro* for activity against *Mycobacterium ulcerans*. Beta-lactams tested were effective alone and displayed a strong synergistic profile in combination with antibiotics currently used to treat BU, i.e. rifampicin and clarithromycin; this activity was further potentiated in the presence of the beta-lactamase inhibitor clavulanate. In addition, quadruple combinations of rifampicin, clarithromycin, clavulanate and beta-lactams resulted in multiplicative reductions in their minimal inhibitory concentration (MIC) values. The MIC of amoxicillin against a panel of clinical isolates decreased more than 200-fold within this quadruple combination. Amoxicillin/clavulanate formulations are readily available with clinical pedigree, low toxicity, and orally and pediatric available; thus, supporting its potential inclusion as a new anti-BU drug in current combination therapies.

## AUTHOR SUMMARY

Buruli ulcer (BU) is a chronic debilitating disease of the skin and soft tissue, mainly affecting children and young adults in tropical regions. Before 2004, the only treatment option was surgery; a major breakthrough was the discovery that BU could be cured in most cases with a standard treatment that involved 8 weeks of combination therapy with rifampicin and streptomycin. However, the use of streptomycin is often associated with severe side effects such as ototoxicity, nephrotoxicity, or hepatotoxicity. More recently, a clinical trial demonstrated equipotency of replacing the injectable streptomycin by the clarithromycin, which is orally available and with fewer side effects. Thus, BU treatment is now moving toward a full orally available treatment of clarithromycin-rifampicin. However, recent studies have reported the emergence of strains resistant to rifampicin and streptomycin. Since no alternatives to rifampin are available, it is considered the cornerstone drug for BU therapy. Strains resistant to clarithromycin could also eventually emerge. Thus, there is a need to find new therapies for BU.

In this work, we describe for the first time the potential inclusion of beta-lactams in BU therapy. More specifically, we propose the use of amoxicillin/clavulanate since it is oral, suitable for the treatment of children, and readily available with a long track record of clinical pedigree. Its inclusion in a triple oral therapy complementing current combinatorial rifampicin-clarithromycin treatment has the potential to counteract resistance development and to reduce length of treatment and time to cure.

## INTRODUCTION

Buruli ulcer (BU), caused by *Mycobacterium ulcerans*, is a chronic debilitating mycobacterial disease of the skin and soft tissue. Although mortality is low, permanent disfigurement and disability is high, mainly affecting children and young adults. BU is found primarily in tropical regions of Africa, South America and the Western Pacific; however, it is also becoming a public health concern in some regions of Australia, especially in the temperate South-Eastern state of Victoria. This worsening epidemic with increased number of cases in the last few years had an estimated cost to Victoria of $2,548,000 in 2016 and ca. $14,000 per patient ^1^.

Before 2004, when the World Health Organization (WHO) published provisional guidance for the management of BU disease ^2^, antibiotics were viewed as relatively ineffective and surgery remained the mainstay of treatment for BU ^3^ Several attempts had been made to develop an effective antimicrobial BU therapy but without desirable outcomes ^4, 5^. In the late 1990s and early 2000s, however, *in vitro* studies demonstrated anti-BU activity of some antibiotics used for the treatment of tuberculosis (TB) and other non-tuberculous mycobacteria, including macrolides ^6^, fluoroquinolones, rifampicin ^7^ and aminoglycosides ^8^. This knowledge led to further *in vivo* studies demonstrating the potential for combining two drugs to provide optimal treatment outcomes ^9–13^. Soon after, clinical evidence demonstrated the effectiveness of a combination of rifampicin plus streptomycin when it was administered for at least 4 weeks ^14^ and that routine implementation of such a therapy was possible in the field ^15, 16^. This game changer therapy involved 8 weeks of daily painful deep intramuscular injections of streptomycin; however, the use of streptomycin is often associated to adverse events such as oto-, nephron- and hepatotoxicy and is restricted in the treatment of pregnant women and young infants. In addition, the lack of an efficacious oral treatment remained one of the main obstacles to decentralizing care at local level in rural areas. Together, these limitations motivated the scientific community to evaluate alternative oral treatments that did not include streptomycin. Clinical studies demonstrated that fluoroquinolones ^17^ or clarithromycin ^18–20^, could also be used in combination with rifampicin. Since clarithromycin is orally available and with fewer side effects compared to the injectable streptomycin, on March 24^th^, 2017, WHO recommended moving toward adoption of a full oral treatment of 8 week daily combination therapy of rifampicin-clarithromycin in its *“Report from the Biennial Meeting of the Buruli ulcer Technical Advisory Group”* ^21^

Although antimicrobial therapy is major step toward the management of this infectious disease, eventual failure due to the development of resistance is anticipated. *M.ulcerans* strains resistant to rifampicin have already been isolated after experimental chemotherapy in mice ^22^ and a recent report described the emergence of *M. ulcerans* strains resistant to rifampicin and streptomycin in the clinic ^23^. WHO currently recommends only four drugs for the treatment of BU: rifampicin, streptomycin, clarithromycin and moxifloxacin ^2^. Rifampicin is the essential partner in combinatorial therapy since its sterilization properties prevents relapse of the disease ^13^, making it the cornerstone drug for BU therapy. As such, no alternatives for rifampicin are currently available and new oral therapeutic options readily available in the clinic should be explored that are suitable for children and pregnant women.

In this study, we are translating knowledge and concepts of drug repurposing and synergies generated in TB R&D programs to assess the potential inclusion of beta-lactams to complement current anti-BU therapy. We proposed the combination of amoxicillin/clavulanate as a new anti-BU treatment in combination with current oral BU therapy, rifampicin and clarithromycin, with the potential of treatment shortening and readily implementation in the field.

## MATERIAL AND METHODS

### Bacterial strains, growth conditions and reagents

*M. ulcerans* strain NCTC 10417 (ATCC Number: 19423; Lot Number: 63210551) was used for initial screening assays. Further validation studies were performed with clinical isolates from different geographical origins: ITM 063846, Benin; ITM 070290, China; ITM 083720 and ITM C05143, Mexico; ITM 941327, ITM C05142 and ITM M000932, Australia; ITM C05150, DR Congo; ITM C08756, Japan, purchased from the Belgian Co-ordinated Collection of Micro-organisms (BCCM).

*M. ulcerans* cells were initially grown at 30°C to an optical density at 600 nm (OD_600_) of 0.5-1.0 in tissue culture flasks containing 7H9 broth supplemented with 0.2% glycerol, 10% OADC and 0.05% (vol/vol) Tyloxapol. Aliquots of 500 μL were then stored at −80°C and the number of colony forming units (CFU) in the freeze stock enumerated. Every experiment was performed starting from a new frozen stock to avoid excessive passage of the original strain. Cells were also routinely passaged on Middlebrook 7H10 agar plates (Difco) supplemented with 10% (vol/vol) OADC to ensure purity of the isolate.

Rifampicin (R3501-1G; Lot Number: SLBH7862V) and meropenem (M2574; Lot Number: 055M4705V) were purchased from Sigma. GlaxoSmithKline provided clarithromycin, streptomycin, clavulanate and all other beta-lactams used in this study.

### Drug susceptibility assays

Minimal Inhibitory Concentrations (MIC) were determined in 7H9 broth supplemented with 0.2% glycerol, 10% OADC and without Tyloxapol using triplicate two-fold serial dilutions of compounds in polystyrene 384- or 96-well plates. MTT [3- (4,5-dimethylthiazol-2-yl)-2,5-diphenyl tetrazolium bromide] was used as the bacterial growth indicator ^24^ For cell density calculations, a culture having an OD_600_ of 0.125 was found to contain approximately 10^7^ cfu/mL. Cultures were sampled (50 μL in 384-well plates or 200 μL in 96-well plates) at a final cell density of 10^6^ cfu/mL and incubated at 30°C in the presence of the drug (or drug combinations) for 6 days before addition of 12.5 μL (384-well plates) or 30 μL (96-well plates) of a MTT / Tween 80 (5 mg/mL / 20%) solution mix. After further overnight incubation at 37°C, OD_580_ was measured. The lowest concentration of drug that inhibited 90% of the MTT color conversion (IC_90_) was used to define MIC values.

### Synergy assays

Checkerboard assays and calculations of the Fractional Inhibitory Concentration Index (FICI) were used to define the degree of pairwise drug interactions, as previously described ^25^. Up to quadruple combinations of rifampicin, clarithromycin, beta-lactams and clavulanate were also tested. For this, checkerboard plates were prepared with rifampicin in the y-axes, the beta-lactam in the x-axes, and clarithromycin added in the z-axes as fixed sub-MIC (1/2, 1/4, and 1/8xMIC values) concentrations for every checkerboard plate; typically the 1/8xMIC plate was used for synergy calculations of the quad combos. When assayed, clavulanate was added at a fixed sub-MIC concentration of 5 μg/mL. Increase efficacy of compounds (synergistic MIC, MIC_syn_)when in combination (fold-MIC reduction) was always reported versus the activity of drugs alone. The Most Optimal Combinatorial Concentration (MOCC) was defined as the lowest possible concentration of every compound that, when assayed together, prevented bacterial growth, i.e. in an isobologram representation this would be the closest point to the axes intersection.

## RESULTS

### Rifampicin has strong synergistic interactions with beta-lactams but not with current anti-BU drugs

Clinically approved beta-lactams representing different sub-families, i.e. meropenem (carbapenems), cephradine and cefdinir (cephems), faropenem (penems) and amoxicillin (penicillins) were assayed *in vitro* in a checkerboard format to assess their synergistic interactions with rifampicin in the absence and presence of clavulanate, a beta-lactamase inhibitor, against the *M. ulcerans* ATCC strain. A pattern of strong synergistic interaction was observed between rifampicin and all the beta-lactam tested (**Figure 1**); however, no interaction was observed when the same assay was conducted using combinations of rifampicin and the currently WHO recommended anti-BU drugs, i.e. streptomycin, clarithromycin and moxifloxacin (**Figure S1**). Dose-response curves indicated that the activity of rifampicin (reflected in MIC reduction) was increased on average 16-32-fold (up to 128-fold in some cases) and, vice versa, the activity of the beta-lactams was strongly enhanced by rifampicin. In the case of amoxicillin, its activity was further increased 512-fold in combination with clavulanate (**Figure S2** and **Table S1**).

**Figure 1.**
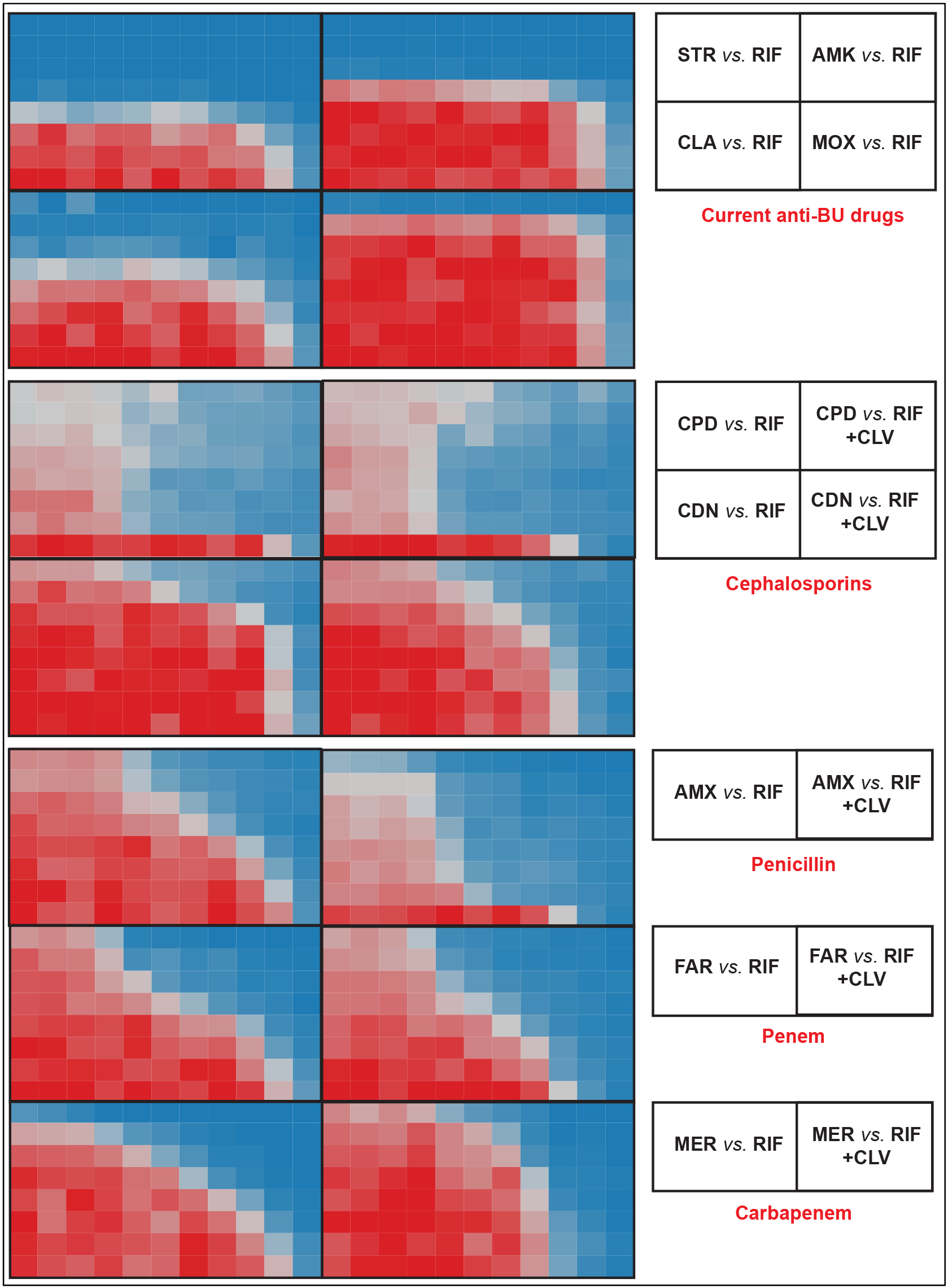
Synergistic profiles of current anti-BU drugs and selected beta-lactams with rifampicin against *M. ulcerans* ATCC 19423. Compounds were assayed in two fold serial dilutions in a checkerboard format. Rifampicin was assayed in the x-axis while other drugs were assayed in the y-axis. RIF, rifampicin; STR, streptomycin; AMK, amikacin; CLA, clarithromycin; MOX, moxifloxacin; CPD, cephradine; CDN, cefdinir; AMX, amoxicillin; FAR, faropenem; MER, meropenem; CLV, clavulanate.

Beta-lactams not only had synergistic interactions with rifampicin but also with clarithromycin, the second drug recommended as first-line anti-BU therapy. These results prompted us to test the inhibitory effect of double clarithromycin-beta lactam, and triple rifampicin-clarithromycin-beta-lactam combinations (Figure 2). Our results indicated that, when in double or triple combinations, much lower sub-inhibitory concentrations were equally potent at inhibiting *M. ulcerans* growth than the additive effects of the compounds alone. MIC values were also lower than in other pairwise combinations. For example, the MIC of amoxicillin was greater than 32 μg/mL; however, its synergistic MIC (MIC_syn_) was reduced to 1 μg/mL in the presence of rifampicin, to 0.25 μg/mL when clavulanate was also added, or to 0.062 μg/mL when both clavulanate and clarithromycin were included together with rifampicin, i.e., an MIC reduction of ca. 500-fold for amoxicillin when in the quadruple combination. Similar results were obtained for combinations of meropenem or faropenem and rifampicin, with MIC reductions as high as 80-fold when tested within triple combinations. In these assays, clarithromycin was added at a fixed concentration of 1/8 its MIC value alone, being its presence critical to achieve the multiplicative effect observe in the quadruple
combinations.

**Figure 2.**
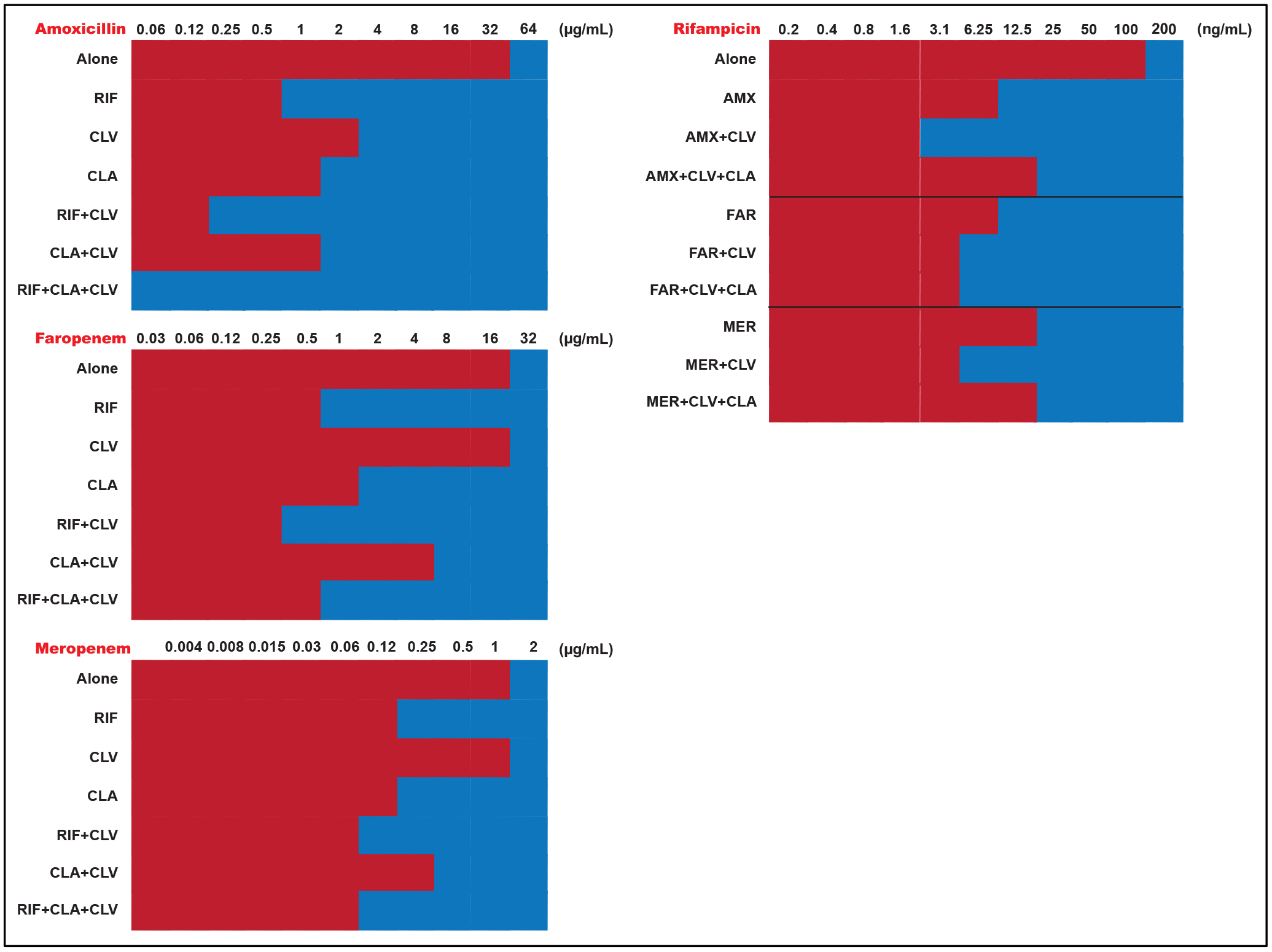
Multiplicative effects of quadruple synergistic combinations including rifampicin, clarithromycin, clavulanate and different beta-lactams against *M. ulcerans* ATCC 19423. The MIC of each compound was compared alone and in the presence of several synergistic combinations at the MOCC (lowest FICI). Clarithromycin is not displayed since it was tested at a fixed 1/8xMIC concentration (MIC_CLA_ = 0.5 μg/mL). Clavulanate was tested at a fixed 5 μg/ml) concentration. AMX, amoxicillin; CLA, clarithromycin; CLV, clavulanate; FAR, faropenem; MER, meropenem; RIF, rifampicin.

### Amoxicillin/clavulanate is highly active against *M. ulcerans* clinical isolates in combination with rifampicin and clarithromycin

Our initial discoveries described above were performed using the reference *M. ulcerans* ATCC strain. In order to validate the pattern of synergistic interactions between rifampicin, clarithromycin and beta-lactams and to gauge the potential of introducing a beta-lactam in the treatment of BU, we expanded our analysis to a collection of *M. ulcerans* clinical isolates from different geographical locations and focused our synergy interaction studies on the amoxicillin/clavulanate combination. When in the quadruple combinations, the activities of all three drugs (clavulanate was added at a fixed 5 μg/mL concentration, more than 20-fold less its MIC) were strongly enhanced; depending on the strain tested, these interactions could range from ca. 5 to 600-fold (rifampicin), ca. 4 to 2,000-fold (amoxicillin) and ca. 20 to 80-fold (clarithromycin) (**Table 1**). Every possible pair-wise and triple combination was also evaluated showing strong synergism between amoxicillin and both rifampicin and clarithromycin, but not between rifampicin and clarithromycin, similar to previously described for the ATCC strain. Clavulanate enhanced the activity of amoxicillin (as expected) but had no effect over clarithromycin and only minor enhancements (in some cases) over rifampicin (**Table S3**).

**Table 1.** Quadruple synergistic combinations among rifampicin, clarithromycin, amoxicillin and clavulanate against *M. ulcerans* clinical isolates.

| Clinical isolate | Geographical origin | MIC (µg/mL) alone |  |  | MICsyn (µg/mL) at the MOCC |  |  | Fold-Reduction at the MOCC |  |  | quad FICI |
| --- | --- | --- | --- | --- | --- | --- | --- | --- | --- | --- | --- |
|  |  | RIF | AMX | CLA | RIF | AMX | CLA | RIF | AMX | CLA |  |
| ITM 063846 | Benin | 0.025 | >32 | 0.125 | 0.0031 | 1 | 0.003 | 8 | 64 | 40 | 0.20 |
| ITM 070290 | China | 0.5 | >32 | 1 | 0.0016 | 0.063 | 0.05 | 313 | 1024 | 20 | 0.09 |
| ITM 83720 | Mexico | 0.1 | 16 | 0.125 | 0.0063 | 0.063 | 0.003 | 16 | 256 | 40 | 0.13 |
| ITM 941327 | Australia | 0.002 | >32 | 0.5 | 0.0004 | 0.031 | 0.025 | 5 | 2065 | 20 | 0.29 |
| ITM C05142 | Australia | 0.1 | 16 | 0.125 | 0.0016 | 0.063 | 0.0031 | 64 | 256 | 40 | 0.08 |
| ITM C05143 | Mexico | 0.05 | 16 | 0.25 | 0.0008 | 4 | 0.0031 | 64 | 4 | 80 | 0.32 |
| ITM C05150 | DR Congo | 0.02 | >32 | 0.25 | 0.0013 | 0.063 | 0.0031 | 16 | 1024 | 80 | 0.12 |
| ITM C08756 | Japan | 0.2 | >32 | 0.25 | 0.0016 | 0.250 | 0.0125 | 128 | 256 | 20 | 0.10 |
| ITM M000932 | Australia | 0.05 | >32 | 0.063 | 0.0001 | 0.125 | 0.0031 | 640 | 256 | 20 | 0.09 |

Clavulanate was added to the quadruple combination at a fixed dose of 5 µg/mL. For FICI calculations, its MIC was considered as 128 µg/mL.
RIF, rifampicin; AMX, amoxicillin; CLA, clarithromycin.
MOCC, Most Optimal Combinatorial Concentration
Fold-Reduction, MIC/MICsyn
FICI ≤0.5 indicates synergy.

Amoxicillin is inactivated by beta-lactamases enzymes, limiting its clinical use. In fact, dose-response studies demonstrated that amoxicillin alone was not active (MIC > 16 μg/mL); however, a strong shift in the dose-response curve was observed by adding clavulanate and this shift was further enhanced when clarithromycin and rifampicin were also present in the combination at sub-MIC concentrations (**Figure 3**), thus confirming our MIC/synergy data and the potential of amoxicillin/clavulanate as a new anti-BU therapy, both alone and in combo with rifampicin, clarithromycin or both.

**Figure 3.**
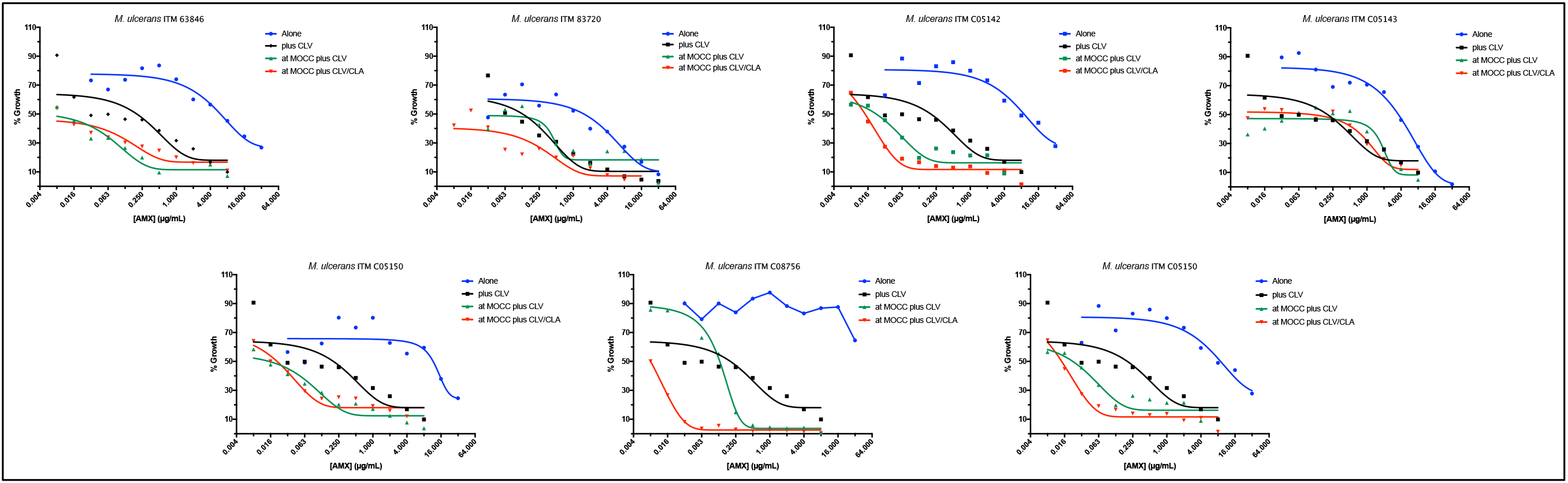
Dose response curves of amoxicillin alone and in combination against *M. ulcerans* clinical isolates. Dose response curves of amoxicillin tested: “alone”, alone; “Plus CLV”, in the presence of clavulanate; “at MOCC plus CLV”, in the presence of clavulanate and rifampicin at the MOCC; “at MOCC plus CLV/CLA”, in the presence of clavulanate and rifampicin and clarithromycin at the MOCC. Clavulanate was tested at a fixed 5 μg/mL concentration. AMX, amoxicillin; CLA, clarithromycin; CLV, clavulanate; FAR, faropenem; MER, meropenem; RIF, rifampicin

## DISCUSSION

In this study, we have *in vitro* characterized the antimicrobial interactions of rifampicin with anti-BU drugs and beta-lactams and found that, while there was no synergy between rifampicin and current anti-BU drugs, it had strong synergistic interactions with beta-lactams. What is more, beta-lactams also displayed synergism with clarithromycin, the other first line drug in BU therapy. Our studies confirmed previous data showing lack of *in vitro* interaction between rifampicin and clarithromycin^26^, thus reinforcing synergy data observed with beta-lactams.

Working with *M. ulcerans* is challenging due to its slow generation time (ca. 48 hours), even slower than *M. tuberculosis.* Current methodologies to perform susceptibility testing against clinical isolates are often time consuming and cumbersome (use of agar proportion methods), thus, requiring several weeks to generate results ^6, 8, 23^.

Improvements have been introduced using luminescent reporter strains ^27^; however, this technology is limited to specific engineered strains and cannot be widespread applied to clinical isolates. Here, we were able to perform synergy studies, obtaining results in just seven days, in a medium-throughput manner against a panel of clinical isolates by using an experimental design previously described for TB research ^25^. Briefly, a synthetic 7H9 mycobacteria broth micro-dilution assay (96- and 384-well plates) was employed coupled to a modification of the MTT method^28^, where the MTT solution was added together with 20% of Tween 80 to solubilize the formazan precipitate in a one-step reaction. We found that MTT provided a more robust signal and wider dynamic range than resazurin-based micro-dilutions assays ^29, 30^ This methodology provides a cost and time effective assay to implement in the clinical practice in drug resistance surveillance campaigns.

Before 2004, surgery was the only option available for the treatment of Buruli ulcer. Over the last decade, therapy has experience a tremendous improvement with the introduction of chemotherapy; however, there is only a limited number of drugs recommended by WHO for BU therapy, namely rifampicin, streptomycin, clarithromycin and moxifloxacin ^2^ Current WHO recommended therapy is a fully oral eight weeks daily regime with a combination of rifampicin and clarithromycin ^20^ Although effective and mostly well tolerated, combination treatment of rifampicin plus streptomycin or clarithromycin are associated with undesirable side effects that might include mild (anorexia, nausea, abdominal pains and altered taste) or severe (deafness, skin rashes, jaundice, shock, purpura or acute renal failure) symptoms ^2^. In such a scenario, where administration of any of these drugs needs to be interrupted, therapeutic options remain limited to the use of moxifloxacin, an antibiotic contraindicated during pregnancy and within the pediatric population. A similar situation would occur in the eventual development of resistance to any of these drugs, especially rifampicin. Since they are administered in pairwise combinations, this would effectively imply monotherapy, further promoting the emergence of resistance. This is similar to already seen in another mycobacterial disease, leprosy, for which this threat was largely ignored and just recently WHO issued guidelines including procedures for the detection of drug resistance ^31^. Thus, an alternative drug regimen would be required to treat resistant *M. ulcerans* strains ^23^.

The WHO recommended regimen allowed, however, cure of small lesions (<5 cm in diameter) without surgery ^15, 18^ Controversy remained regarding the best approach for large lesions (>10 cm) whether to combine prolonged chemotherapy (up to 12 weeks) with surgery ^32^ or to delay this intervention ^33^, since postponing surgery reduced both the duration of hospitalization and wound dressing. It would be thus desirable to develop a new therapy that would reduce both duration of treatment and time to healing after therapy completion. Reducing the duration of therapy from 8 weeks to 6, or even fewer weeks, would be a major breakthrough in the management of BU. Before this ambitious goal can be achieved, intermittent drug administration has been proposed as a strategy to facilitate treatment supervision in the field by health care workers. Along this line, experiments using mice models of infection suggested the use of rifapentine, a rifampicin analog with longer half-life, instead of rifampicin that has to be administered daily; theoretically allowing for a fully oral intermittent regimen (with clarithromycin) against *M. ulcerans* ^34^ Although promising, experience from TB research has revealed inconsistencies between animal model data and clinical predictability. A mouse study demonstrating that daily dosing of rifapentine would lead to cure of TB in three months or less in the standard regimen, led to the Tuberculosis Trials Consortium Study 29 (TBTC 29) of 531 patients; however, in TBTC 29, daily rifapentine in patients was no better than daily rifampin ^35^ Moreover, such a BU regimen would still rely on a two-drug combination therapy, with all the shortcomings above described.

The history of TB chemotherapy teaches us that combination therapy is critical for optimal cure outcome. The first anti-TB drug, streptomycin, discovered in 1944, was immediately used in monotherapy for treatment of TB patients. After an initial improvement over the first months, health deterioration ensued due the development of resistant *M. tuberculosis* strains, rendering streptomycin ineffective. To overcome these limitations the first combination therapy for the treatment of a disease was developed using para-aminosalicylic acid together with streptomycin. However, this therapy still required 18 months of treatment. It was only after the discovery of rifampicin in 1965 and the subsequent introduction of this drug in the combination therapy that treatment could be reduced to 9 months. Today, the WHO recommended TB treatment include a combination of four drugs for 2 months (rifampicin, isoniazid, ethambutol, and pyrazinamide) plus a continuation phase of 4 months with just rifampicin and isoniazid ^36^ Translating this knowledge into BU therapy, it remains clear that more drugs need to be added to the current rifampicin-clarithromycin combination in order to improve and shorten the duration of treatment.

Rifampin is the cornerstone drug for TB (and BU) therapy. There is a direct relation between dose increase and therapy efficacy ^37^ due to its bactericidal and sterilizing activity in a dose-dependent manner ^38^. However, the current WHO recommended dose of 10 mg/kg (current 600 mg daily) is not used routinely at its optimal clinical dose ^39^ In fact, it was introduced in 1971 based on pharmacokinetic, toxicity, and cost considerations but some recent studies suggest that rifampicin dose could be safely increased to 35 mg/kg daily^37^. Follow up efficacy studies revealed that only doses of 35 mg/kg ^40^, but not lower ^41^, had a bacteriological effect on time to culture conversion, suggesting that synergistic partners could serve to improve rifampicin efficacy without compromising tolerability and toxicity.

Drug development for neglected diseases such as BU, mainly affecting developing countries, is especially complicated due to lack of interest from the main scientific and industrial communities and, as a consequence, lack of investment. To overcome these limitations and speed up the discovery and development process, we applied knowledge gathered in TB R&D programs using a drug repurposing approach ^24, 25, 42, 43^ In these programs, we aimed to improve the activity of rifampicin within a synergistic combination with the hypothesis that if rifampicin efficacy could be increased, TB therapy could be shortened, while reducing toxicity. In the course of this program we (and others ^44^) identify that beta-lactams strongly increased the bactericidal and sterilizing properties of rifampicin ^25^

Beta-lactams are one of the largest groups of antibiotics available today with an exceptional record of clinical safety in humans. With over 34 FDA-approved beta-lactams, they are the most widely used antibiotics in history constituting ~50% of all antibiotic prescriptions worldwide ^45^ Used for decades to treat diseases caused by bacteria, they had been traditionally considered to be ineffective for the treatment of mycobacterial infections (mainly TB) due to the presence of a beta-lactamase (BlaC) able to degrade these antibiotics, and the hydrophobic nature of the mycobacterial cell envelope that limits beta-lactam access to their cellular targets ^46^. However, after a seminal publication describing the *in vitro* activity of the meropenem plus clavulanate combination against multi-drug (MDR) and extensively drug resistant (XDR) strains of *M. tuberculosis* ^47^ and anecdotally use in salvage therapies for XDR patients ^48^, the first study convincingly demonstrating the clinical efficacy of beta-lactams was recently published ^43^ These studies provided evidence of their anti-mycobacterial clinical potential alone and in synergistic combinations with rifampicin, opening a new avenue to identify new drugs and optimize current BU therapy.

Most beta-lactams tested in this study were active against *M. ulcerans* and enhanced the anti-BU activity of rifampicin to different degrees. Cephradine is a first-generation cephalosporins developed in the 1960s, also recently described to be active *in vitro* against *M. tuberculosis* ^25^; however, cephradine was long ago discontinued and access to other first-generation cephalosporins, such as cefadroxil or cephalexin, is limited in many countries. Cefdinir is a third-generation cephalosporin active against pneumonia, skin and soft tissue infections, although with low oral absorption ^49^. It is currently used in the clinic, widely distributed and access to it could be readily available; however, its synergistic profile with rifampicin was weaker compared to other beta-lactams (**Table S1**). Although meropenem was active against pulmonary TB in a recent clinical study ^43^, it needs to be administered intravenously, not a practical approach in under-resourced countries where oral drugs are required. Faropenem, an orally administered beta-lactam, did not show activity in the same clinical trial due to the low drug exposure in plasma after oral administration ^43^. Finally, amoxicillin/clavulanate showed good activity and very strong synergistic interaction with rifampicin.

The combination of amoxicillin plus clavulanate is a broad-spectrum antibacterial available for clinical use in a wide range of indications and is now used primarily in the treatment of community-acquired respiratory tract infections ^50^. It was first launched in the UK in 1981; by the end of 2002, it was clinically available as various formulations in over 150 countries around the world. In addition to high efficacy, it has a well-known safety and tolerance profile, including for pregnancy and paediatric used, based on over 819 million patient courses worldwide, with the main contraindication being allergy to penicillin derivatives ^51^.

For TB treatment, amoxicillin/clavulanate is included in Group 5 (anti-TB drugs with limited data on efficacy and long-term safety in the treatment of drug-resistant TB) of the WHO 2011 TB drugs classification and in Group D3 (add-on agents, not core MDR-TB regimen components) of the WHO 2016 MDR-TB drugs classification ^52^. In 1983, Cynamon *et al.* reported the *in vitro* bactericidal activity of amoxicillin/clavulanate against 15 isolates of *M. tuberculosis,* at concentrations of amoxicillin lower than 4 μg/mL ^53^, and some years later, Nadler *et al.* case reported the effective treatment of MDR-TB patients with the addition of amoxicillin/clavulanate to the second-line therapy ^54^ Two contradictory follow up 2-days Early Bactericidal Activity (EBA) clinical studies suggested that its activity was comparable to that reported for anti-TB agents, other than isoniazid ^55^, but also questioned its role in the treatment of tuberculosis ^56^ The dosing interval of the amoxicillin/clavulanate therapy might explain these differences; while in the first EBA it was divided into three daily doses, it was given as a single high dose in the second one. More recently, a 14-days EBA study demonstrated activity of a combination of meropenem plus amoxicillin/clavulanate ^43^; it remains to be determined whether this activity was due to any of the components alone or the combination therapy as a whole^57^. In fact, *in vitro* studies have demonstrated synergistic interactions among amoxicillin and meropenem (among other beta-lactams), rifampicin and ethambutol against *M. tuberculosis* ^25, 57–59^

In our *in vitro* assays with *M. ulcerans* clinical isolates, we found that amoxicillin had no activity (typically MIC values > 16 μg/mL) but that its MIC could be reduced to 1 μg/mL in the presence of clavulanate (**Table S3**), similar to previously reported to the closely related *M. marinum* species and other non-tuberculosis mycobacteria ^60^ It also displayed strong synergistic interactions with rifampicin and clarithromycin and, in quadruple combinations, its activity was enhanced up to 2,000-fold in some cases, with average MIC ranges between 0.031 to 0.25 μg/mL (**Table 1**). For infections caused by other bacterial pathogens, susceptibility breakpoints of amoxicillin/clavulanate are established at ≤ 2 μg/mL (or ≤ 4 μg/mL for high-dose formulations) and mean peak plasma concentrations of amoxicillin range from 7.2 to 17 μg/mL, depending on the formulation ^51^, well above the synergistic MIC values reported in this work. Thus, according to our *in vitro* data, amoxicillin/clavulanate could have an important role in the treatment of BU alone and, more importantly, in combination with current first-line anti-BU therapy since no pharmacological drug-drug interactions are described among amoxicillin/clavulanate and rifampicin or clarithromycin ^50^.

The bacteriological efficacy of penicillins is dependent on the time its free plasma concentrations remain above the MIC (time over the MIC value, fT>MIC). For other bacterial infections, it has been estimated that a fT>MIC of ca. 30-40% of the dosing interval is required for bactericidal activity ^61^. In the case of *M. ulcerans,* this target therapy could be achieved using standard amoxicillin/clavulanate formulations of 500/125 mg (4:1) or 875/125 mg (7:1) administered three times a day, or the high-dose extended release formulation of 2000/125 (16:1) that would allow administration twice a day ^51^, an important consideration for treatment compliance in under-resourced settings.

But, what could be the benefit of adding amoxicillin/clavulanate to the current anti-BU therapy? Besides being able to treat secondary infections associated with BU lesions, it has been proposed that the median time to healing is related to the bacterial load in the lesions at the beginning of therapy and the presence of persister bacteria^62^; in fact, healing of up to two thirds of patients occurs within 25 weeks from the start of treatment but for some patients this can take up to a year. One of the reasons for this slow healing could be due to a high initial bacterial load. In fact, active infection late into the recommended 8-week course of antibiotic therapy could be found in slowly healing lesions ^32, 63^. Amoxicillin/clavulanate could have the potential to address these issues since it is extremely effective at targeting extracellular bacteria with a rapid bactericidal activity and extensive histopathologic studies demonstrate that *M. ulcerans* is essentially confined in extracellular areas of necrosis in skin ^6^. Amoxicillin/clavulanate could have an important role at rapidly reducing the initial bacterial load. In addition, *in vitro* studies have demonstrated the sterilizing activity of synergistic combinations of beta-lactams and rifampicin ^25^, which could target those persistent populations, thus reducing treatment shortening and healing times. Rapid bacterial killing would also imply a reduction in the risk of development of resistance, as already reported for rifampicin and streptomycin ^23^; however, even in the scenario of infections caused by bacteria resistant to rifampicin, this could still be re-introduced for BU therapy if it was administered with amoxicillin/clavulanate, as previously demonstrated in *M. tuberculosis* ^25^. Finally, because of their synergistic interactions with clarithromycin, it could replace rifampicin in the treatment of HIV patients under anti-retroviral therapy.

Our study comes as well with some limitations. Although representative of different geographical origins, we only tested one ATCC strain and 9 clinical isolates. Further *in vitro* studies with a larger set of *M. ulcerans* clinical isolates would be needed to assess the full clinical potential and coverage of a triple combination including rifampicin, clarithromycin and amoxicillin/clavulanate. We did not test the combinations using *in vivo* models of *M. ulcerans* infection. Mice are a sub-optimal *in vivo* model to evaluate the activity of beta-lactams since their pharmacokinetics and efficacy in mice do not predict those found in humans ^64^; this is in part due to the fact that mice express an enzyme that degrades beta lactams (renal dehydropeptidase I, DPH-I) at levels that are several orders of magnitude higher than in humans ^65, 66^, thus effectively reducing the time beta-lactams are over the MIC value. In addition, the translational power of such *in vivo* mice models have been questioned in recent TB clinical trials that were based on mice data ^35, 67^. Because amoxicillin/clavulanate is a well-known antimicrobial with a clear track record of safety over decades of use we believe that direct evaluation in clinical trials would be the fastest route to improve treatment of BU patients; however, questions remain on posology and length of treatment and additional dose optimization PKPD studies would be needed. To such an end, the hollow fiber system has proven a useful tool in the TB field, recently endorsed by the European Medicines Agency^68^

## CONCLUSIONS

By using a repurposing approach and using technology already developed in TB R&D programs, we have identified amoxicillin/clavulanate as a new potential anti-BU drug to be used alone and in combination therapy with rifampicin and clarithromycin, current first-line anti-BU drugs, to reduce the length of therapy and the time to healing. Disruption of the gut microbiota is the main associated side effect of long-term use of amoxicillin/clavulanate, mainly caused by the presence of clavulanic acid in the formulation. Based on the strong synergistic interactions among amoxicillin with rifampicin and clarithromycin, amoxicillin alone might be added to the full course of a shorter therapy. However, because the main role of amoxicillin/clavulanate in the anti-BU therapy would be to reduce the initial bacterial load found in the lesions, we propose the use of high-dose extended release formulations during just the first two weeks of therapy.

## ACKNOWLEDGMENTS

We would like to thank Drs. Santiago Ferrer, David Barros and Lluis Ballell (GlaxoSmithKline) for their continuous support to the successful outcome of this work, and the Biology Unit of the GlaxoSmithKline TB DPU for technical support. We are also grateful to Begoña Gracia (University of Zaragoza) for excellent laboratory and strain management and to Frangoise Portaels and Miriam Eddyani (Institute of Tropical Medicine, ITM, Antwerp) for providing the *M. ulcerans* clinical isolates.

This work was supported by grants from a People Programme (Marie Skłodowska Curie Actions) of the European Union’s Seventh Framework Programme (FP7/2007-2013) under REA agreement No. 291799 (Tres Cantos Open Lab Foundation - COFUND programme), from the European Union’s Horizon 2020 research and innovation programme under the Marie Skłodowska-Curie grant agreement No. 749058, and from the Tres Cantos Open Lab Foundation to S.R.-G.

## AUTHOR CONTRIBUTIONS

Conceived the study: A.M-L. and S.R-G.; Designed the experiments: M.P.A-C., and S.R-G.; Performed experiments: M.P.A-C., R.G.d.R., A.L.Q. and S.R-G.; Analyzed data: M.P.A-C., and S.R-G; Critically reviewed the manuscript: C.J.T and A.M-L.;Wrote the manuscript: S.R-G. All authors approved the final version of the paper.

## ADDITIONAL INFORMATION

Supplementary information accompanies this paper: Tables S1, S2 and S3, and Figures S1 and S2.

Competing financial interests: R.G.d.R., and A.M.-L. are employees of GlaxoSmithKline, a producer of the generic drug amoxicillin/clavulanate. All other authors declare no conflicts of interest. The funders had no role in the study design, data collection and interpretation, or the decision to submit the work for publication.

## SUPPLEMENTARY INFORMATION

**Table S1.**
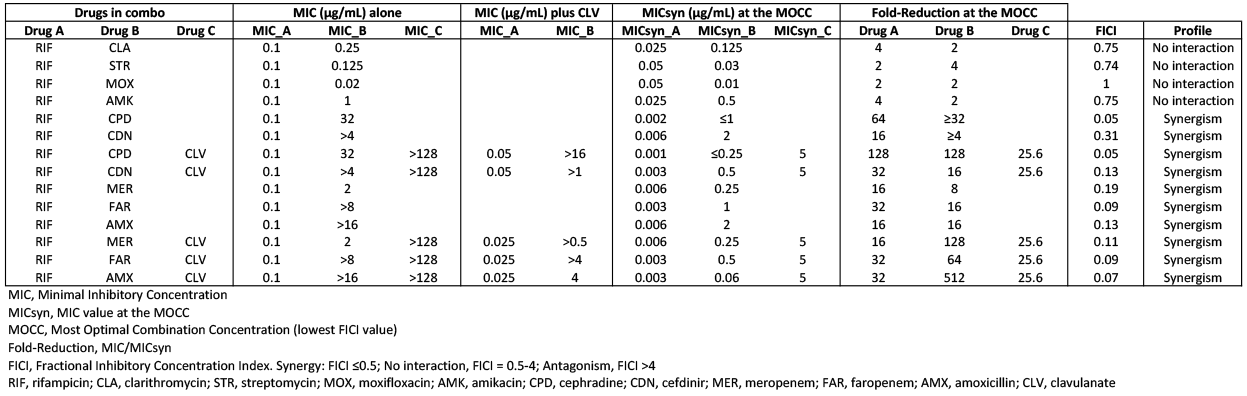
Synergistic interactions among rifampicin and beta-lactams against *M. ulcerans* ATCC 19423. Data supporting Figure 1.

**Table S2.**
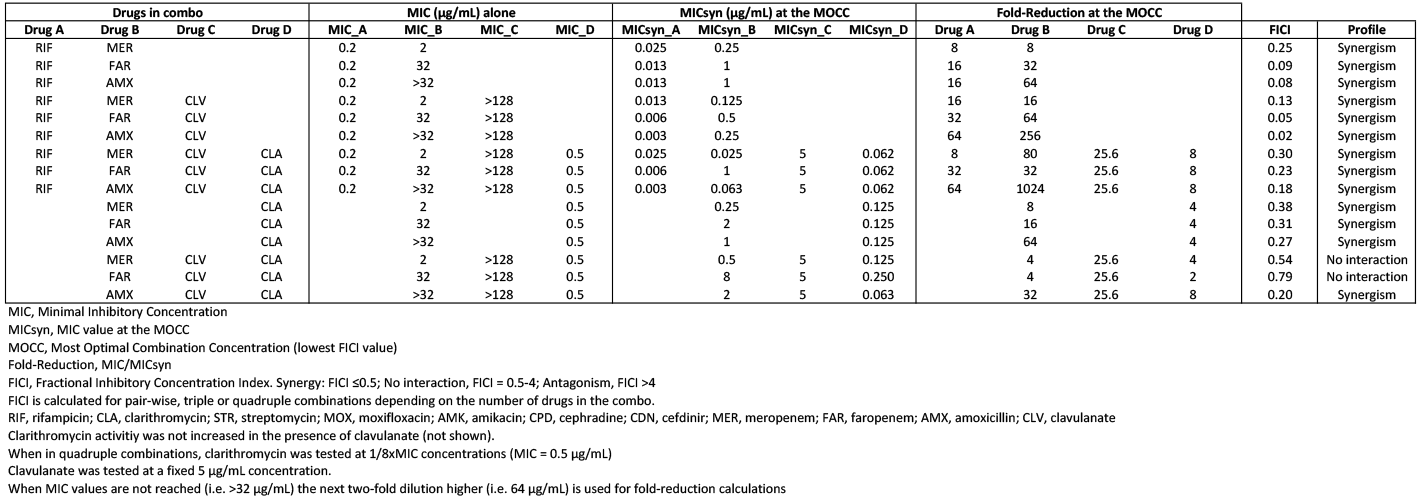
Quadruple synergistic combinations among rifampicin, clarithromycin, beta-lactams and clavulanate against *M. ulcerans* ATCC 19423. Data supporting Figure 2.

**Table S3.**
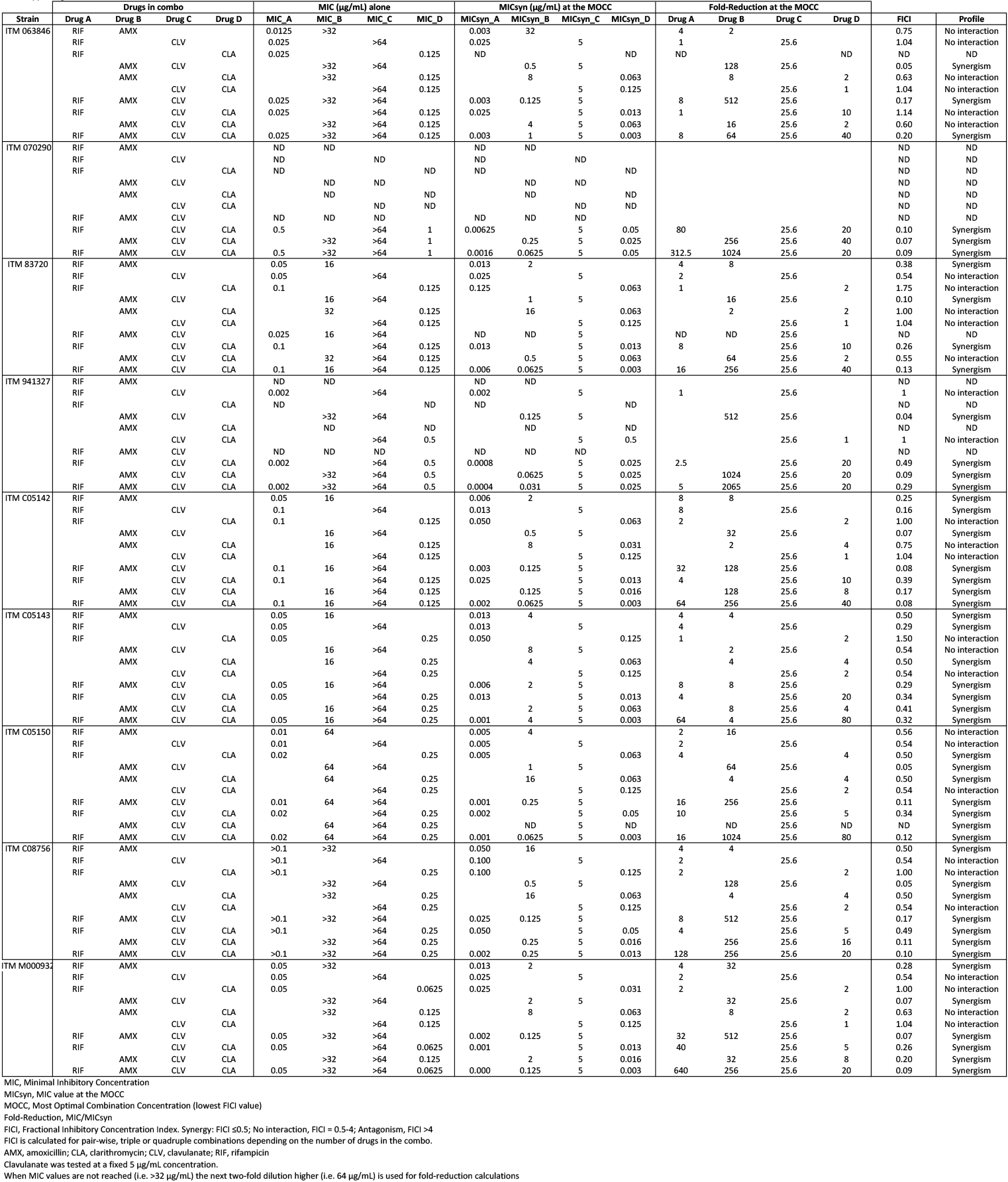
Quadruple synergistic combinations among rifampicin, clarithromycin, beta-lactams and clavulanate against *M. ulcerans* clinical isolates. Data supporting Table 1.

**Figure S1.**
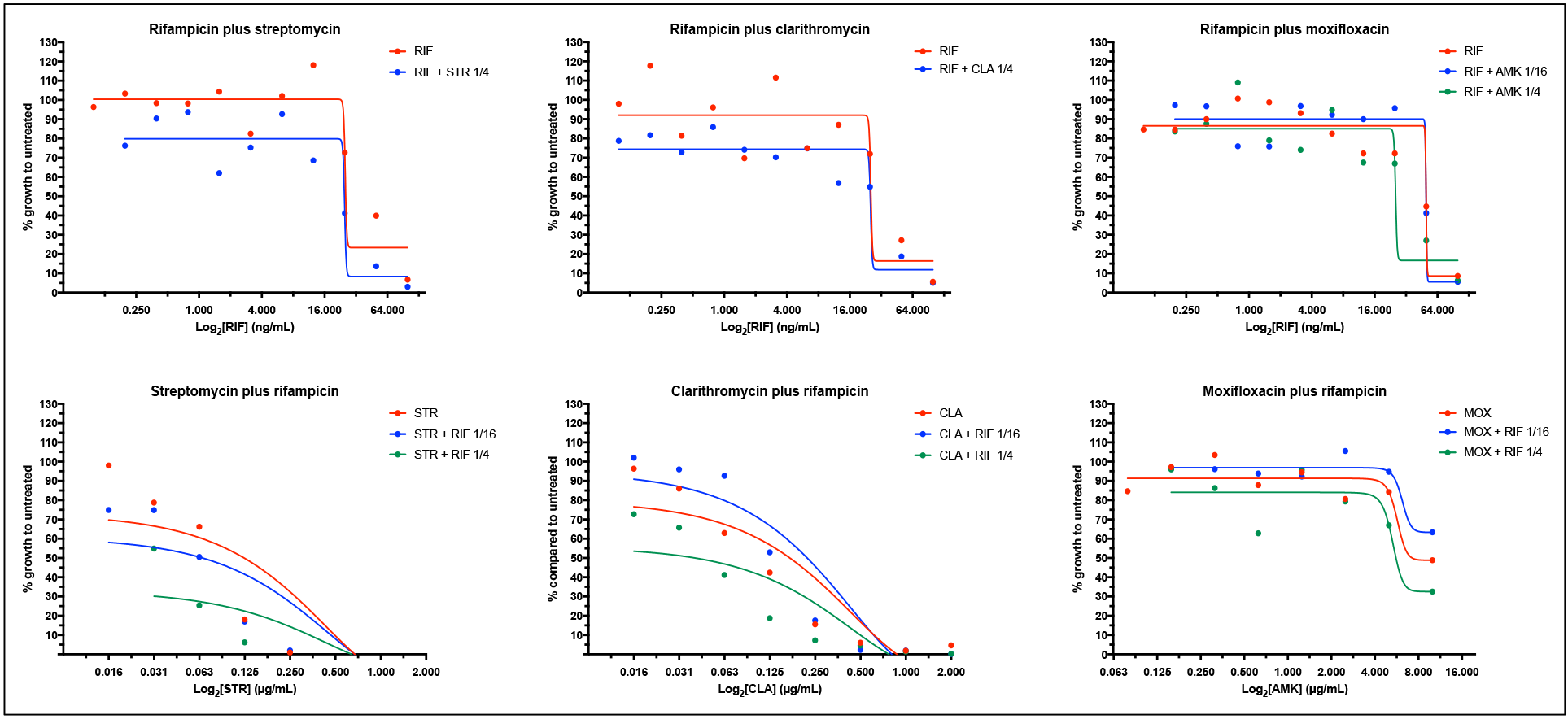
Dose response curves of rifampicin in the presence of anti-BU drugs against *M. ulcerans* ATCC 19423. Dose response curves plotted are those of the drug alone and in the presence of l/4×MIC and l/16×MIC concentrations of the synergistic partner. RIF, rifampicin; CLA, clarithromycin; MOX, moxifloxacin; STR, streptomycin

**Figure S2.**
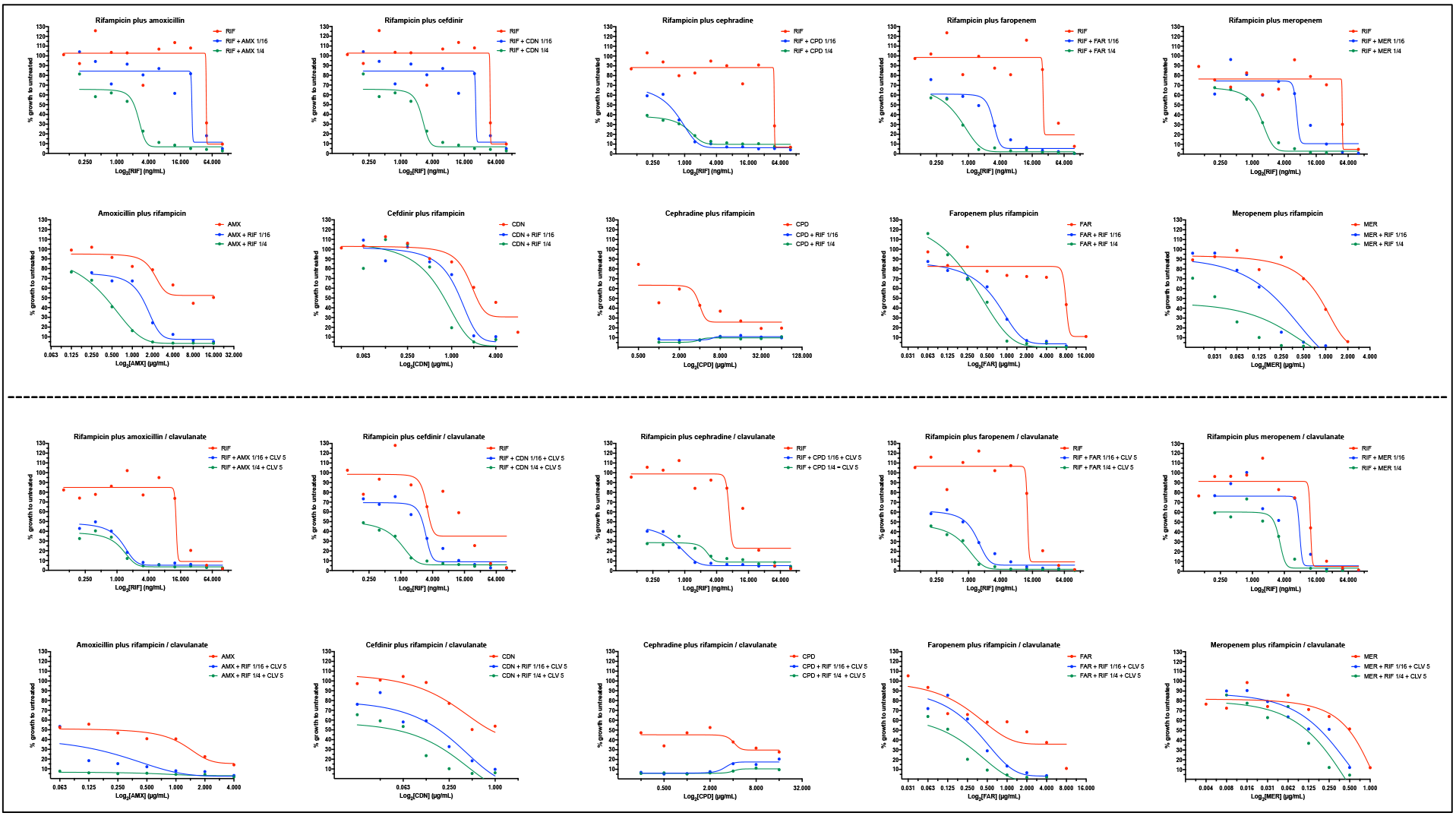
Dose response curves of rifampicin in the presence of beta-lactams, and viceversa, without (top panel) and with clavulanate (bottom panel) against M. ulcerans ATCC 19423. Dose response curves plotted are those of the drug alone and in the presence of l/4×MIC and l/16×MIC concentrations of the synergistic partner. Clavulanate was added at a fixed concentration of 5 μg/mL. RIF, rifampicin; AMX, amoxicillin; CDN, cefdinir; CPD, cephradine; FAR, faropenem; MER, meropenem

